# Pathway diversity: a resilience metric sensitive to agency applied to a lake eutrophication problem

**DOI:** 10.1101/2025.06.09.658728

**Authors:** Vitor Hirata Sanches, Joseph Guillaume, Takuya Iwanaga, Sarah Clement, Steven J. Lade

## Abstract

Measuring resilience has been a longstanding challenge in social-ecological research. While there are established resilience metrics, they often do not account for agency, which is crucial in social-ecological systems. Pathway diversity is a recent approach to measuring resilience that integrates systems thinking with individual-based elements, such as agency. According to this approach, resilience is larger if actors have more decision pathways and can maintain them over time. Here we measure resilience using pathway diversity within a lake eutrophication model to analyse (a) the extent to which it is sensitive to changes in agency and (b) its compatibility with more established resilience metrics. Our findings reveal that pathway diversity is sensitive to regime shifts and can provide early warnings of them. Through five policy decision-making scenarios, we then show how pathway diversity can capture constraints on decision makers’ capability and agency that affect the system’s resilience. We find that higher pathway diversity is associated with greater capability of decision-makers to influence the system. Pathway diversity addresses a critical gap by incorporating agency into a resilience metric while remaining compatible with established metrics. This work shows the potential of pathway diversity to identify resilience-based policy implications.

## 1. Introduction

Resilience is an increasingly popular concept in academia and policy (Reyers et al., 2022), and multiple approaches have been developed to measure it (Dakos and Kéfi, 2022; Krakovská et al., 2023; Polain de Waroux et al., 2024; Sanches et al., 2025). Social-ecological resilience, the view used here, defines resilience as the capacity of a system to respond to change, including the capacity to persist, adapt and transform (Folke, 2006; Walker et al., 2004). Rooted in ecological resilience, the concept has evolved through interaction with community, development, social, engineering, disaster, and psychological resilience (Baggio et al., 2015; Quinlan et al., 2016). Its interdisciplinary nature and broad possible conceptualisations make it particularly challenging to measure (Allen et al., 2019; Baggio et al., 2015; Quinlan et al., 2016). Common approaches to measuring aspects of social-ecological resilience include compound indicators (Sharifi, 2016; Laurien et al., 2022) and time-based performance indicators (Steinmann et al., 2024), among others (González-Quintero and Avila-Foucat, 2019; Polain de Waroux et al., 2024; Sanches et al., 2025; Steinmann et al., 2024).

Pathway diversity is a recent approach to measuring resilience by integrating system and agency perspectives. According to pathway diversity, a system is more resilient if actors have more sequences of actions available, i.e., pathways, over a certain time horizon (Lade et al., 2020). The approach can capture agentic dimensions and agency by focusing on options available to agents. At the same time, it captures system properties such as feedback by considering the effect of choosing one option on future option availability. The approach builds on the widely recognised importance of diversity for resilience and the idea that response and functional diversity are key mechanisms for promoting resilience (Biggs et al., 2015; Elmqvist et al., 2003; Walker et al., 2023). A key benefit of pathway diversity is its ability to capture both system and agent-based properties (Lade et al., 2020). However, the effects of changes in agency on pathway diversity are not fully demonstrated in a case study and being a new approach, it is not clear if it is compatible with more established and well-known resilience metrics.

In this study, we measure resilience using pathway diversity in a lake eutrophication model to analyse the extent to which pathway diversity is sensitive to changes in agency and whether it is compatible with more established resilience metrics. Having a quantitative resilience metric that is sensitive to agency addresses key gaps identified in previous studies (Cote and Nightingale, 2012; Otto et al., 2020; Sanches et al., 2025), while ensuring consistency with established resilience metrics and theories is essential for integrating a new approach into the resilience literature. Lake eutrophication is one of the first and best documented cases of regime shifts, which are a large and persistent change in the system structure and function (Biggs et al., 2018; Folke, 2006), in this case, from a state with moderate plant production and clear water to one with high plant production and murky water (Carpenter et al., 1999). The lake eutrophication model proposed by Carpenter et al. (1999) is foundational in resilience research (Dakos and Kéfi, 2022; Scheffer et al., 2001), and over time it has been used in multiple decision-making analyses (Peterson et al., 2003; Singh et al., 2015; Quinn et al., 2017; Rougé et al., 2013).

We compare pathway diversity with two ecological resilience metrics: distance to the basin threshold and early-warning signals metrics. We choose ecological resilience metrics because social-ecological resilience is rooted in ecological resilience and these metrics can capture properties that are relevant for analysing the lake eutrophication problem like regime shifts and feedbacks. The two selected metrics are among the most well-known and established in ecological resilience (Dakos and Kéfi, 2022; Krakovská et al., 2023). Metrics based on the shape of potential landscape compute the potential landscape to analyse system stability and are one of the earliest approaches to analysing resilience (Holling, 1973; Walker et al., 2004). A very common metric in this class is the distance to the basin threshold, which measures how far the current state is from the basin threshold. It measures how strong a disturbance must be to cause a regime shift and indicates the likelihood of a regime shift. Early-warning signals is a model-free approach to analyse the system’s stability based on statistical patterns in the time series of the system state (Dakos et al., 2012; Scheffer et al., 2009). For example, systems usually show high variance close to regime shifts, and there are multiple statistical measures that can assess anomalies as signs of proximity to critical transition (Dakos and Kéfi, 2022; Scheffer et al., 2009).

The importance of agency to social-ecological resilience is widely recognised, yet few resilience metrics are sensitive to agency (Brown & Westaway, 2011; Cote and Nightingale, 2012; Sanches et al., 2025). Resilience metrics derived from ecological approaches naturally tend to overlook agency and how actors influence social-ecological systems (Cote and Nightingale, 2012; Olsson et al., 2015; Sanches et al., 2025). An agent’s capability to take action (agency), its motivation and external systemic constraints to action became more important as the literature shifted from ecological resilience to social-ecological resilience (Cote and Nightingale, 2012; Hahn and Nykvist, 2017). Agency is particularly relevant to building adaptive and transformative capacities (Bohle et al., 2009; Brown & Westaway, 2011; Haider and Cleaver, 2023; Westley et al., 2013). Debates on ‘intentional’ or ‘guided’ transformations and the need to detect and anticipate thresholds and regime shifts in policy implicitly or explicitly underscore the importance of agency (Jozaei et al., 2022; Morgan et al., 2024). Studies argue that understanding agents’ motivations and practices, as well as external factors affecting their decision-making, such as power and governance structures, are essential to understanding how they respond to change (Brown & Westaway, 2011; Cote and Nightingale, 2012; Haider and Cleaver, 2023). Despite the evidence on the importance of agency for resilience and progress on qualitative resilience assessments in including agency, few resilience metrics are sensitive to agency, and researchers have called for more studies incorporating agency into resilience metrics and models (Otto et al., 2020; Sanches et al., 2025).

## 2. Methods

### 2.1. The model

We analyse the resilience of a lake to eutrophication using a dynamical system model (Carpenter et al., 1999). Eutrophication is a phenomenon that occurs when excessive nutrient inputs, typically from agriculture, drive a lake from a regime characterised by low to moderate levels of plant production and relatively clear water into a new regime characterised by high plant production and murky water (Carpenter et al., 1999). We consider a hypothetical town where decision-makers must choose how much phosphorus from agricultural run-off is allowed to enter a shallow lake. The amount of phosphorus in the water column (*x*) is determined by Equation 1 and depends on four terms: (1) the anthropogenic influx of phosphorus, mainly produced by agricultural activities; (2) the removal of phosphorus caused by the sedimentation, outflow and plant consumption of phosphorus in the water column; (3) the recycling of phosphorus, representing sediments and phosphorus consumers that released back phosphorus to the water column; (4) a stochastic process that follows a log-normal distribution, representing natural uncontrolled influx of phosphorus. To simplify the analysis, the last term was only used in the early-warning signal simulation, where stochasticity is required for the method to function.

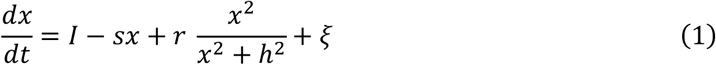

where *I* is the input of phosphorus per time, our decision variable; *s* is the rate of phosphorus loss per time; *r* is the maximum rate of recycling; *h* is the recycling half-saturation rate, the amount of phosphorus where the recycling reaches half of its maximum; and *ξ* is the natural influx of phosphorus, a stochastic process that follows a log-normal distribution with mean *μ* and standard deviation *σ* (*LN*(*µ, σ*)) at each time *t*, where *t* is measured in years.

The recycling term introduces a non-linearity in the system and can cause bi-stability depending on the parameters used. The recycling term has a sigmoidal shape, meaning the phosphorus added by the recycling term suddenly increases upon crossing a certain phosphorus threshold. The shape of this term is based on empirical experiments and is related to the amount of oxygen in the lake (Carpenter et al., 1999). For an intermediate influx of phosphorus, our decision variable, the system will have two stable fixed points (Figure 1), one below and one above the recycling rate phosphorus threshold. Depending on the initial condition position relative to the unstable equilibrium (red line), the system will be attracted either to the eutrophicated (polluted) equilibrium (blue line) or the oligotrophic (clean) equilibrium (green line). At low influx levels, only the clean equilibrium exists, and at high influx levels, only the eutrophicated equilibrium persists (Figure 1).

**Figure 1:**
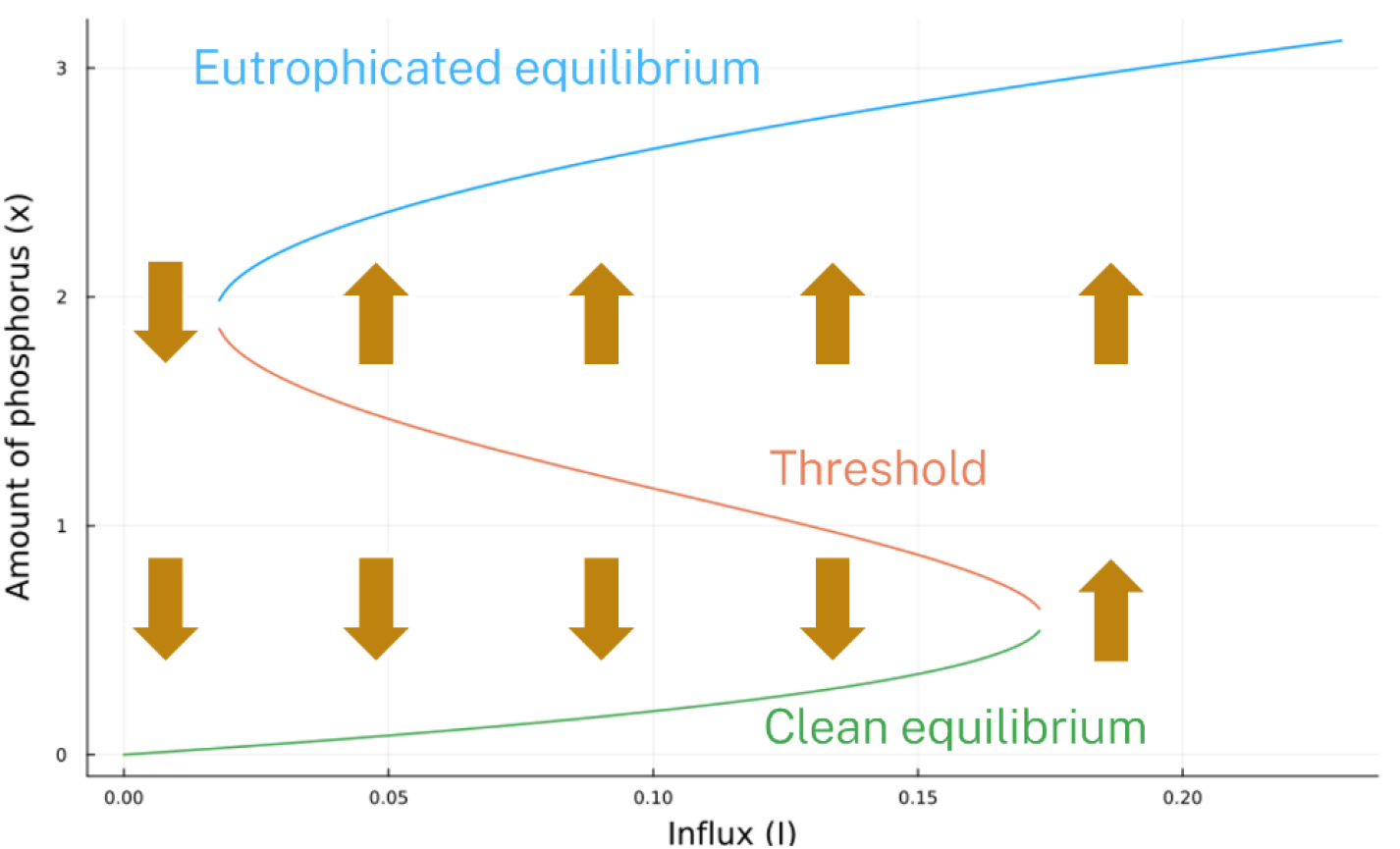
Bifurcation diagram showing the system’s equilibrium as a function of the influx of phosphorus. The parameters used here and throughout the paper are *I*=variable, *s*=0.65, *r*=2.5, *h*=1.95, *μ*=0.1, *σ*=0.01 following the parametrisation done in Dakos and Kéfi (2022). The vertical arrows indicate the direction the system will move. When the system starts above the threshold, it moves toward the eutrophicated (polluted) equilibrium, and when it begins below the threshold, it moves toward the oligotrophic (clean) equilibrium.

### 2.2. The implementation of pathway diversity

To compute pathway diversity, we simulate the lake eutrophication model following all decision pathways up to a time horizon. The decision variable for this model is the input of phosphorus per time (*I*), which town-level regulations can control. We assume that polluting activities follow the maximum allowable influx of phosphorus set by the rules. While this affects the interpretation of the results and limits the generalisability of this study, this assumption is consistent with previous uses of the lake eutrophication problem. The first step in computing pathway diversity is determining the options available for a particular initial condition (Figure 2a). We start by discretising the decision space (*I*) into equally spaced intervals and select the first *n* options excluding zero (Figure 2a, bottom). We exclude *I* = 0 because this is hard to enforce, but we explore the possibility of a very low influx under Section 2.4. To compute the number of available options (*n*), we assume that when the lake is more polluted, the decision-maker has fewer options available and can only choose options with lower phosphorus input per time, therefore, enforcing more restrictive environmental regulations. We implemented this feature by defining a function *n*: ℝ → ℕ that determines the number of options available for a given amount of phosphorus (*x*) (see Figure 2a, middle, for the plot of the function):

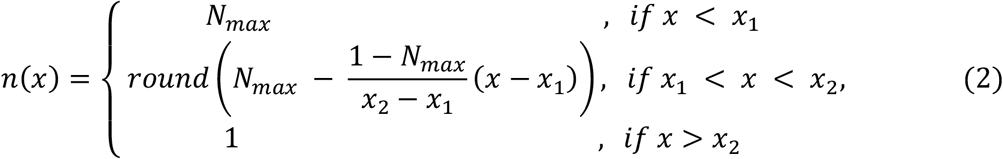

where *N*_*max*_ is the maximum number of options, and *x*_1_ and *x*_2_ are parameters that determine the region where n(x) varies. We fixed *N*_*max*_ = 10 due to computational limitations, *x*_1_ = 0.4 to represent a lower threshold where the system will always be in the clean basin of attraction (Figure 1) and *x*_2_ = 3.0 to represent an extreme upper threshold.

**Figure 2:**
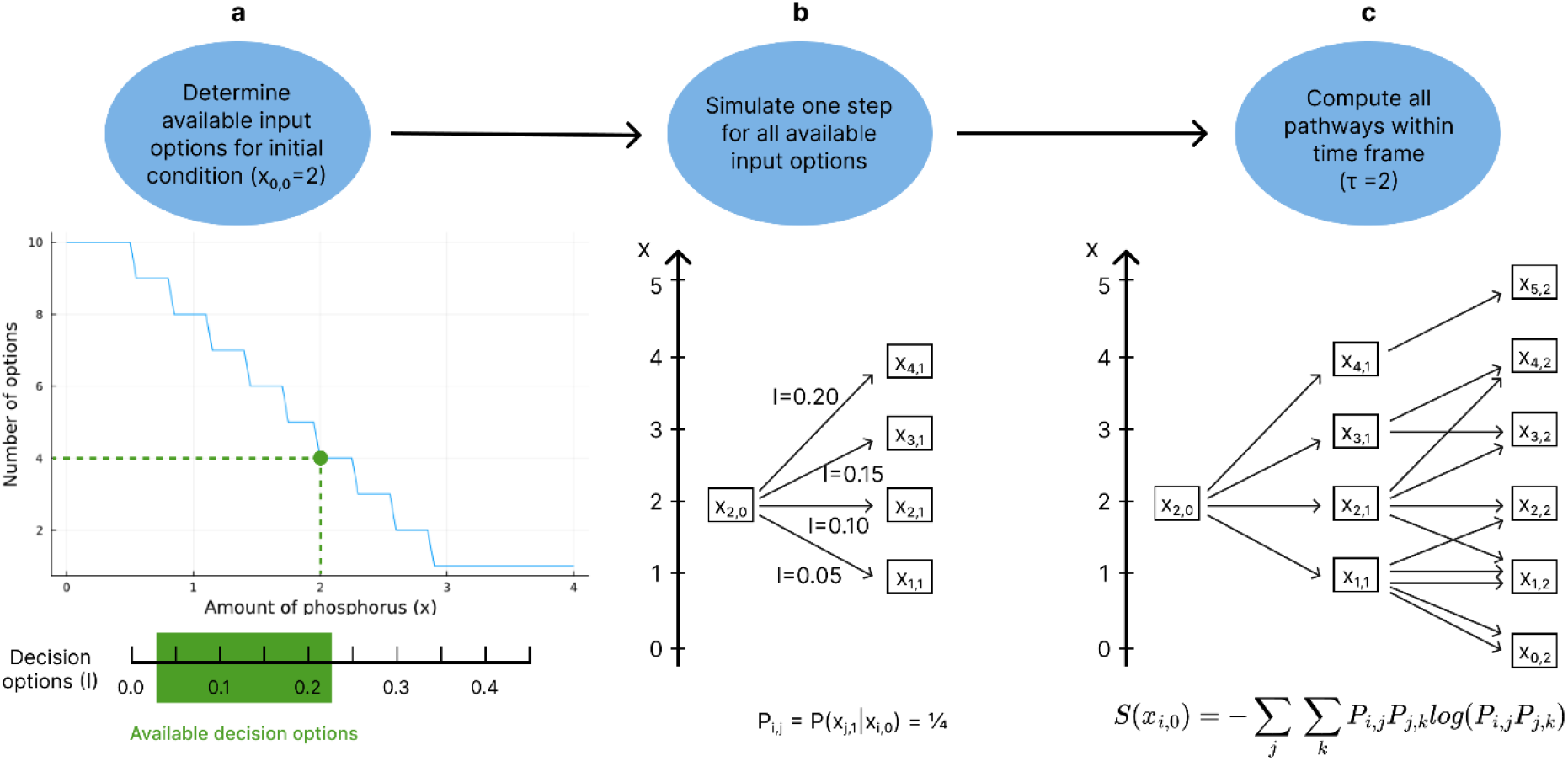
Steps to implement pathway diversity in an example with an initial amount of phosphorus x=2. Step 1 (a): Compute the number of available options (n) for the initial condition. The available options (green area) will be the first n options in the discretised decision space except for *I=0*. In the example, *x=2* and *n(x)=4*, so four options will be available: 0.05, 0.10, 0.15 and 0.20. Step 2 (b): Simulate a time step for all the available options. All options have the same probability of being chosen. In this case, four options are available, so each will have a probability of 1/4. Step 3 (c): Simulate all time steps considering all options available up to the time horizon. To simplify, we chose a time horizon of two decisions. The pathway diversity will be computed with Equation 2 using the probability of starting at *x*_2,0_ and following each pathway *P*_*i,j*_*P*_*j,k*_. The influx of phosphorus values and the number of pathways are illustrative and do not necessarily represent values used in the model.

The second step in computing pathway diversity is to simulate one decision step of the model (Figure 2b). We perform simulations of the system for all available options with a fixed time-step between decisions, five years by default. We assume the probability of choosing each option is the same for all the options available at the step (Figure 2b). This simplification is valid in highly uncertain situations where no option is more desirable than the other. This was used as a default scenario, and we explored alternative methods to define this probability (see Section 2.4).

The final step is to repeat the previous step until the time horizon is reached and then aggregate the results to compute pathway diversity (Figure 2c). Repeating the previous step, we iterate through each pathway, take its final phosphorus amount and simulate one more decision step for all available options. We stop the simulation and compute pathway diversity upon reaching the model time horizon. Following previous definitions, pathway diversity is calculated using causal entropy (Lade et al., 2020; Wissner-Gross and Freer, 2013), which uses the probability of following each pathway from start to end:

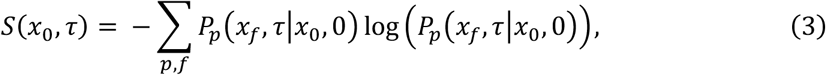

where *P*_*p*_(*x*_*f*_, *τ*|*x*_0_, 0) is the conditional probability of the system transitioning from phosphorus amount *x*_0_ in time 0 to *x*_*f*_ in time τ through the pathway *p*. The sum will be made over all possible pathways *p* to all possible final phosphorus amount *x*_*f*_ the system can reach.

The maximum pathway diversity will happen when the system always has *N*_*max*_ options at all decisions, and the probability of all decisions is equal (higher evenness). In this case, the pathway diversity for a given number of decisions (*n*_*d*_) is

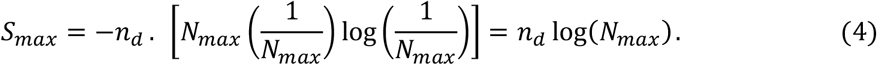

All our results show the pathway diversity normalised by the maximum pathway diversity to facilitate comparisons over different scenarios and parameterisations.

### Comparison between Pathway Diversity and Other Ecological Resilience Metrics

To analyse the relationship between pathway diversity and distance to the basin threshold, we assess whether pathway diversity changes depending on which basin the system starts in and how far it is from the threshold. We compute pathway diversity at multiple initial states throughout the state space (amount of phosphorus). Since pathway diversity is designed to capture system-level dynamics (Lade et al., 2020), we expect it to be sensitive to the regime shift by exhibiting two distinct regimes in its values: one typical of the clean basin of attraction and another of the eutrophicated basin. Accordingly, we expect pathway diversity to gradually change with distance to the basin threshold and show its most significant change upon crossing the threshold region. To test this expectation, we compute the derivative of pathway diversity with respect to the state. If the biggest change in pathway diversity happens in the threshold region, we should observe a peak in its derivative near the threshold. Because the threshold depends on the phosphorus influx (Figure 1), we do not expect a sharp peak, but rather a broad one around the threshold region. To reduce fluctuations caused by the discretisation of the decision space, we apply a LOESS filter before computing the derivative.

To make the early warning signal analysis, we simulate a situation with a linear increase of influx of phosphorus, where the system slowly approaches the regime shift. We assume this situation serves as the initial condition to the pathway diversity analysis, but does not influence it, i.e. when computing pathway diversity, we follow the same process defined in Section 2.2, and all options can be chosen. We use two of the most common indicators of early-warning signals: variance and autocorrelation (Dakos et al., 2012). We follow a standard application of the method, detrending the original time series with a LOESS filter to reduce the noise, applying the two metrics on the detrended series with a rolling window of around half of the time series length and applying the Kendall-τ statistic to find the significance of the early warning signal (Dakos et al., 2012). The Kendall-τ statistic is a rank correlation index that indicates the strength of a trend. To apply this metric, we examine the residuals and select the time step where the signal deviates from a zero mean. We truncate all time series at this point before computing the Kendall-τ statistic to avoid using artificially inflated data points. To assess whether the pathway diversity time series has a significant trend and can indicate an early warning signal, we follow a similar approach and apply the Kendall-τ statistic. However, since pathway diversity is not a statistical metric in the same sense as variance or autocorrelation, we do not use the detrended series or apply a rolling window.

### 2.4. Scenarios for constraints on agency

To analyse how constraints on agency impact pathway diversity, we simulate five policy decision-making scenarios (Table 1; Figure 3). We conceive agency as the capability of decision-makers to take actions, conditioned by internal values and desires and by structural factors such as power, legitimacy (i.e., support or acceptance by others) and institutional constraints (Brown and Westaway, 2011; Duncan, 2019). Through the scenarios, we explore whether pathway diversity can capture how the policy decision-making context influences agency by enabling or constraining the ability of decision-makers to make choices and act on them.

**Table 1.** Description of five policy decision-making scenarios and their implementation.

| Scenario | Policy decision-making context | Implementation |
| --- | --- | --- |
| 1. Reduced decisions | Institutional rigidity makes it harder to change and implement new policy | Reduced frequency of decisions, i.e., taking decisions every 15 years instead of 5 years |
| 2. Allow restrictive option | A stronger government can bear the political costs of an unpopular decision between farmers | The lowest possible influx of phosphorus value is decreased and is close to banning influx. |
| 3. Favour lower influx | Both the government and popular interest groups share the desire for more conservative environmental options | The probability of choosing an option is higher for a lower influx of phosphorus. |
| 4. Minimal change | Opposing interest group pressures make radical changes in policy harder | The probability of choosing an option is proportional to the distance to the current option. |
| 5. Reactive decisions | Following a crisis, the government adopts a reactive, crisis-driven approach to policymaking. | The probability of choosing an option depends on the last change in the amount of phosphorus —decreases favour high influx, increases favour low influx. |

**Figure 3:**
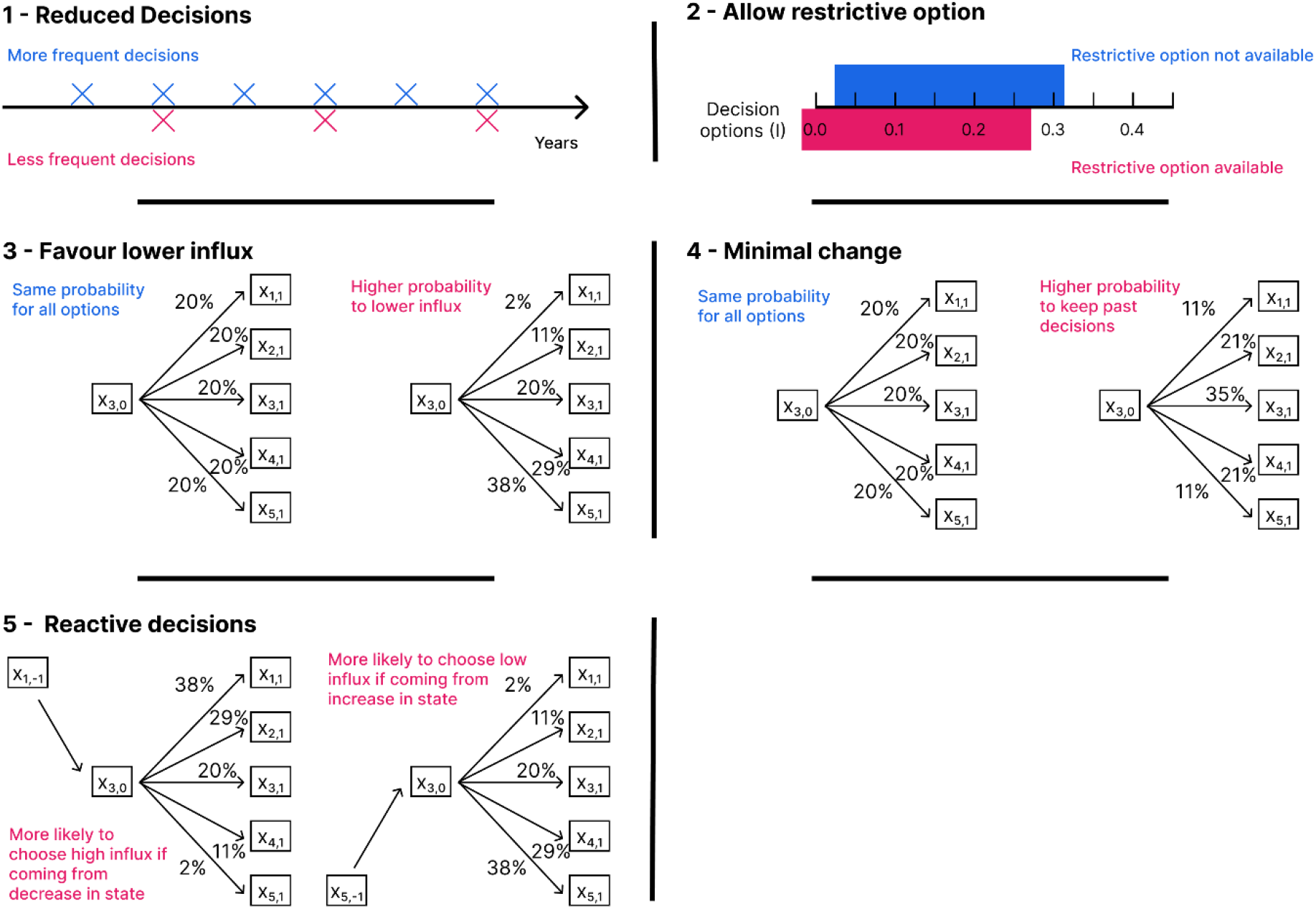
Illustration of five scenarios used in the model. Each box defines one scenario by showing changes made in the scenario (red) compared to the default scenario (blue). The numbers chosen are similar to the ones used on the model but are illustrative, see Section 2.4 for more information.

Each scenario represents a simplified policy decision-making context that captures diverse institutional and policy influences on decision-making (Table 1). In the *reduced decisions* scenario, institutional rigidity (Carpenter and Brock, 2008; Enqvist et al., 2016) constrains agency by limiting how frequently decisions are made. In the *allow restrictive option* scenario, stronger community support increases the government’s legitimacy (Cosens, 2013) and shifts the space of feasible decisions. In the *favour lower influx* scenario, alignment between the decision-maker and community values (Jones et al., 2016) fosters a shared preference for environmentally conservative choices. In the *minimal change* scenario, a rigidity trap caused by opposing interest groups reinforces path dependence (Enqvist et al., 2016; Méndez et al., 2022) and limits change. In the *reactive decisions* scenario, a crisis context constrains agency by pressuring decision-making and resulting in short-term, crisis-driven policies marked by high path dependence (Enqvist et al., 2016; Méndez et al., 2022).

Each scenario changes a decision-making parameter to cover diverse changes in the pathway diversity methodology (Figure 3). We vary the frequency of decisions (scenario 1), the decision space (scenario 2) and how the probability of choosing an option is determined (scenarios 3 to 5). These scenarios can also be interpreted as a sensitivity analysis of the pathway diversity parameters. In the *reduced decisions* scenario, we decrease the decision frequency from once every 5 years to once every 15 years. In the *allow restrictive option* scenario, we shift the decision space from [0.04, 0.30] to [0.02, 0.30], keeping the same maximum number of options and increasing the influx step value. For scenarios 3, 4, and 5, we assign initial weights to each option as described in the following sentences and then normalise the probabilities so that the sum over all options equals to one. In the *favour lower influx* scenario, we assign higher weights to lower influx options using the weighs *w*_*q*_ = 4(*n* − *q*) + 1, where *q* = 1, 2, …, *n* is one option (e.g., [13, 9, 5, 1] for *n=4*). In the *minimal change* scenario, we assume the probabilities have weight *w*_*q*,r_ = 13 − 4*d*(*q*, r), *d* ≤ 3, and *w*_*q,r*_ = 0, *d*(*q, r*) > 3, where *r* is the past option and *d*(*q*, r) = |*q* − r| is the option distance (e.g. [1, 5, 9, 13, 9, 5, 1, 0], for *r*=4, *n*=8). In the *reactive decisions* scenario, we assume the probabilities have weight *w*_*q*_ = 4(*q* − 1) + 1, Δ*x* < −0.2, *w*_*q*_ = 1, = −0.2 < Δ*x* < 0.2, *w*_*q*_ = 4(*n* − *q*) + 1, 0.2 < Δ*x* < 0.8,*w*_*q*_ = 10(*n* − *q*) + 1, Δ*x* > 0.8, where Δ*x* is the past change in the amount of phosphorus (e.g., [13, 9, 5, 1] for *n=4 and* Δ*x* = 0.4 and [1, 5, 9, 13] for *n=4 and* Δ*x* = −0.4).

## 3. Results

We show how pathway diversity compares with distance to the basin threshold in Section 3.1 and early-warning signal metrics in Section 3.2. We show how pathway diversity behaves when policy decision-making context constrain or enable the decision-maker agency in Section 3.3.

### 3.1. Pathway diversity and distance to the basin threshold

Pathway diversity is sensitive to the system threshold, being higher in the clean basin of attraction and lower in the eutrophicated basin of attraction (Figure 4a). When the system starts in the clean basin of attraction, most pathways lead to states in the same basin of attraction, where more pathways are available over time, and thus pathway diversity is higher (Figure 4a). When the system starts in the eutrophicated basin of attraction, the amount of phosphorus almost entirely remains locked in in the same basin of attraction, where fewer pathways are available, and thus pathway diversity is lower (Figure 4a). In both cases, the higher the distance to the basin threshold, the stronger the effect of increasing or decreasing pathway diversity is. The threshold point varies depending on the influx of phosphorus (Figure 1), and in the threshold region, we observe a transient behaviour between the clean and eutrophicated basin of attraction.

**Figure 4:**
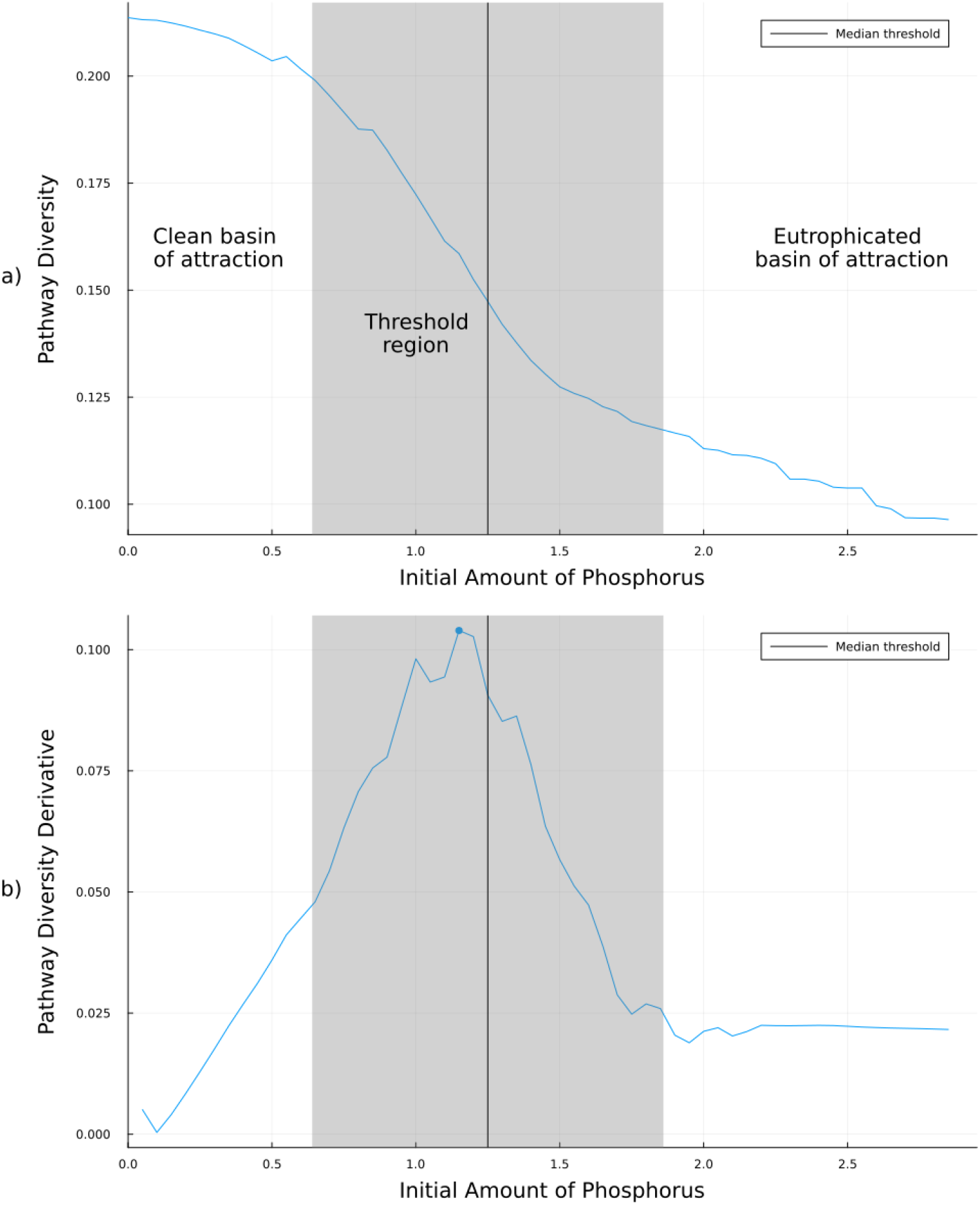
Pathway diversity for different initial amounts of phosphorus (a) and the derivative of that curve (b). The threshold position varies - depending on the influx - within the range shown in grey and has a median indicated by the black vertical line. In b), we show the derivative of a smoothed version of the pathway diversity. The blue point indicates the peak on the derivative, which happens in the threshold region. For this figure, we simulated a time horizon of 35 years.

We observe a peak in the pathway diversity derivative in the threshold region, indicating that the most significant change in pathway diversity happens in this region (Figure 4b). We previously expected the regime shift to be the main factor influencing pathway diversity because the shift causes fundamental changes in the system dynamics which pathway diversity is designed to capture. Thus, Figure 4b confirms that pathway diversity not only captures the regime shift but that the shift is responsible for its most significant change. Additionally, we note that the derivative of pathway diversity reaches higher values for longer time horizons (Appendix S1: Figure S1), that is, when the system is simulated for longer times. This result indicates that both the decline in pathway diversity and its sensitivity to the regime shift become more pronounced as the pathway diversity time horizon increases.

We can further validate the number of pathways leading to different states depending on the initial condition by looking at the state probability distribution over time (Figure 5). We note that lower initial states lead to higher variance in the distribution; that is, most of the probability remains in the clean basin of attraction, but a smaller quantity goes to the eutrophicated basin of attraction (Figure 5a-b). On the other hand, higher initial states lead to a more constrained distribution and pathways remain locked in the eutrophicated basin of attraction (Figure 5c-d).

**Figure 5:**
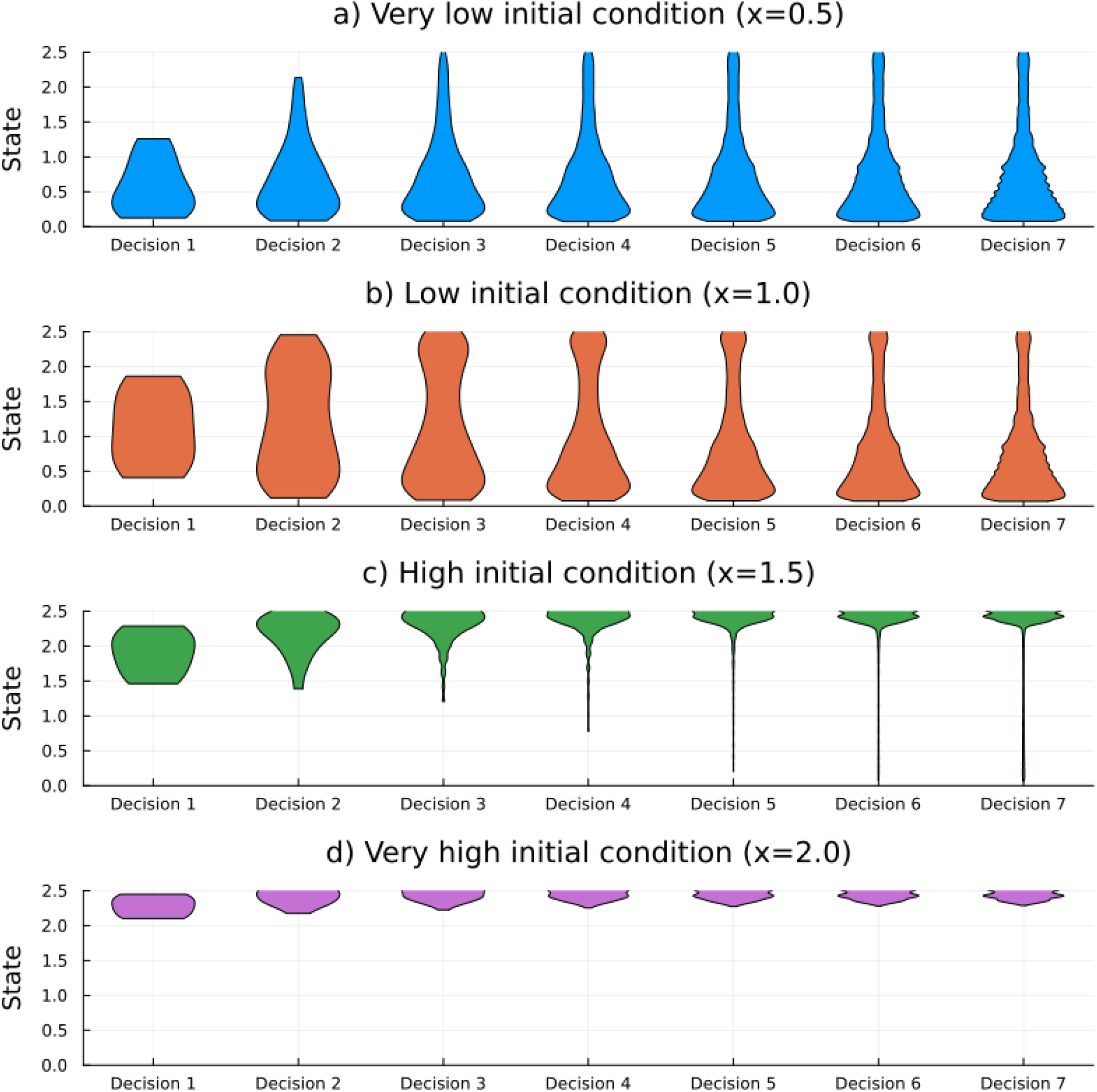
State (*x*) probability distributions for different initial conditions (*x*(0)). The initial condition (amount of phosphorus) varies such that the system starts in (a) the clean basin of attraction, (b) the left of the threshold region, (c) the right of the threshold region, and (d) the eutrophicated basin of attraction. Lower initial conditions lead to distributions with a higher probability of states remaining in the clean basin of attraction and a lower probability of states being in the eutrophicated basin (a, b). Higher initial conditions gradually increase lock-in effects, with the probability distribution becoming constrained within the eutrophicated basin of attraction (c, d).

### 3.2. Pathway diversity and early warning signals metrics

Pathway diversity provides early indications of proximity to regime shifts. Close to a regime shift, pathway diversity declines as more pathways cross the basin threshold and get locked in at a region with fewer options and pathways available (Figure 6). The decline in pathway diversity can be detected using the Kendall-τ statistic, which yields values similar to those obtained from the variance and autocorrelation time series. Therefore, decreasing pathway diversity can indicate an approaching regime shift, like increasing variance and autocorrelation.

**Figure 6:**
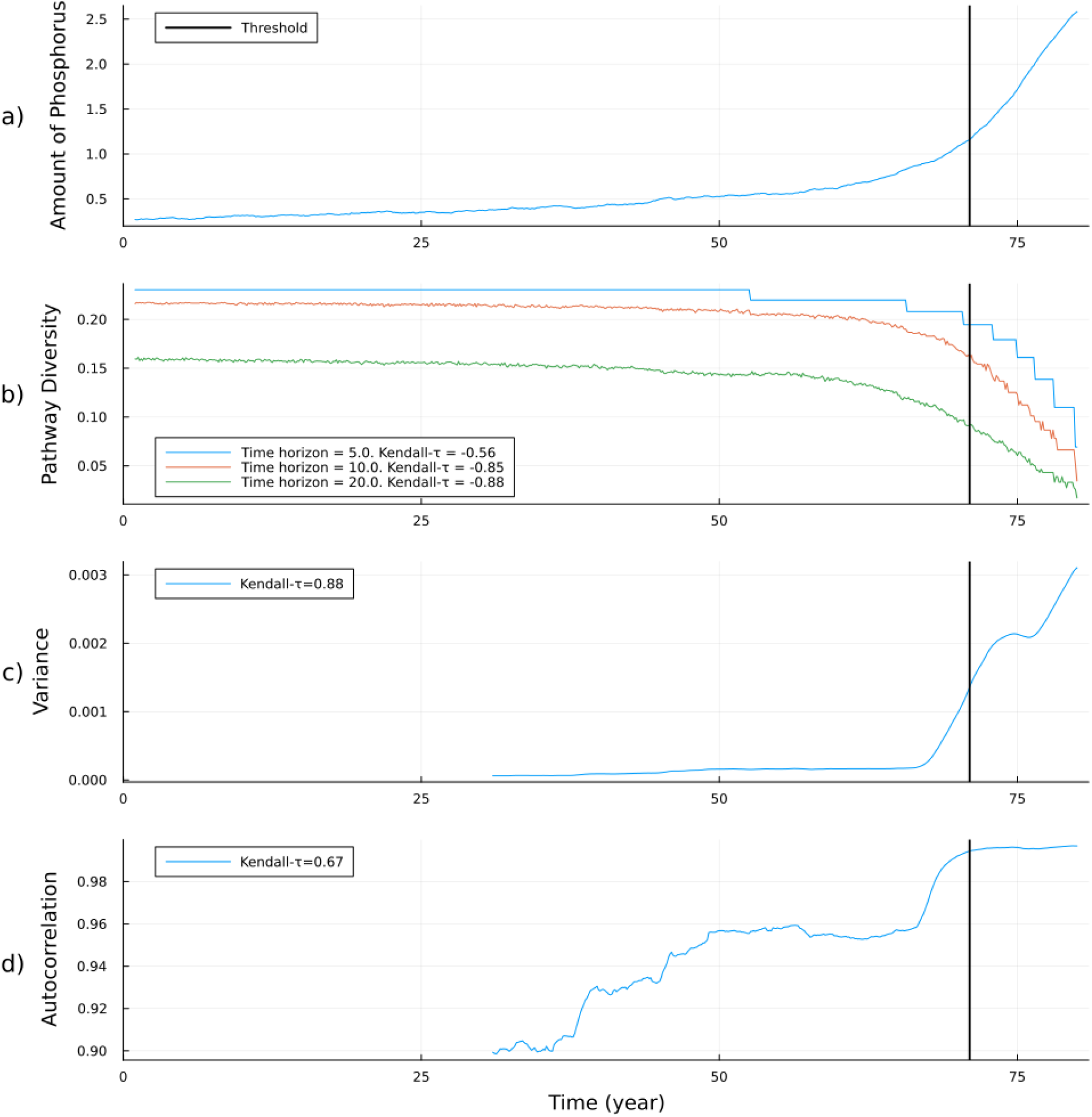
Comparison of pathway diversity and early warning signals. (a) Simulated amount of phosphorous (x), with a linear increase in the influx of phosphorus per time and white noise. Time series of pathway diversity (b) and two early-warning signal metrics, variance (c) and autocorrelation (d). The Kendall-τ was computed at the year 65, because at this time step, the residuals start to deviate from the mean value of zero (Appendix S1: Figure S2).

The trend of declining pathway diversity is stronger at longer time horizons, as indicated by a Kendall-τ statistic further from zero (Figure 6b). For longer time horizons, pathway diversity appears to decrease earlier, likely because the sensitivity to the initial state is higher before the threshold. In this case, a small change in state can lead to a larger proportional shift in the number of pathways that enter and become locked into the eutrophicated basin of attraction.

Early-warning signal metrics and pathway diversity rely on different input data and have different use cases. In this study, pathway diversity is computed using a dynamical system model, incorporating information about how the system evolves under varying parameters. In contrast, early-warning signal metrics are based solely on time series. Pathway diversity requires well-developed models or expert input to estimate the probabilities of different decisions, whereas early-warning signals are model-free and can be applied directly to observed data.

### 3.3. Pathway diversity and change of agency

Our results show that pathway diversity can capture how factors constraining or enabling agency influence resilience. We use five policy decision-making scenarios that affect the decision-maker’s capability to take action (agency). A scenario can represent a decision-maker with higher or lower capability to make decisions. In general, having more agency, in the sense of fewer constraints on decision-making, leads to more pathway diversity when compared to the default scenario, though the effect varies for different regions of the state space (Figure 7).

**Figure 7:**
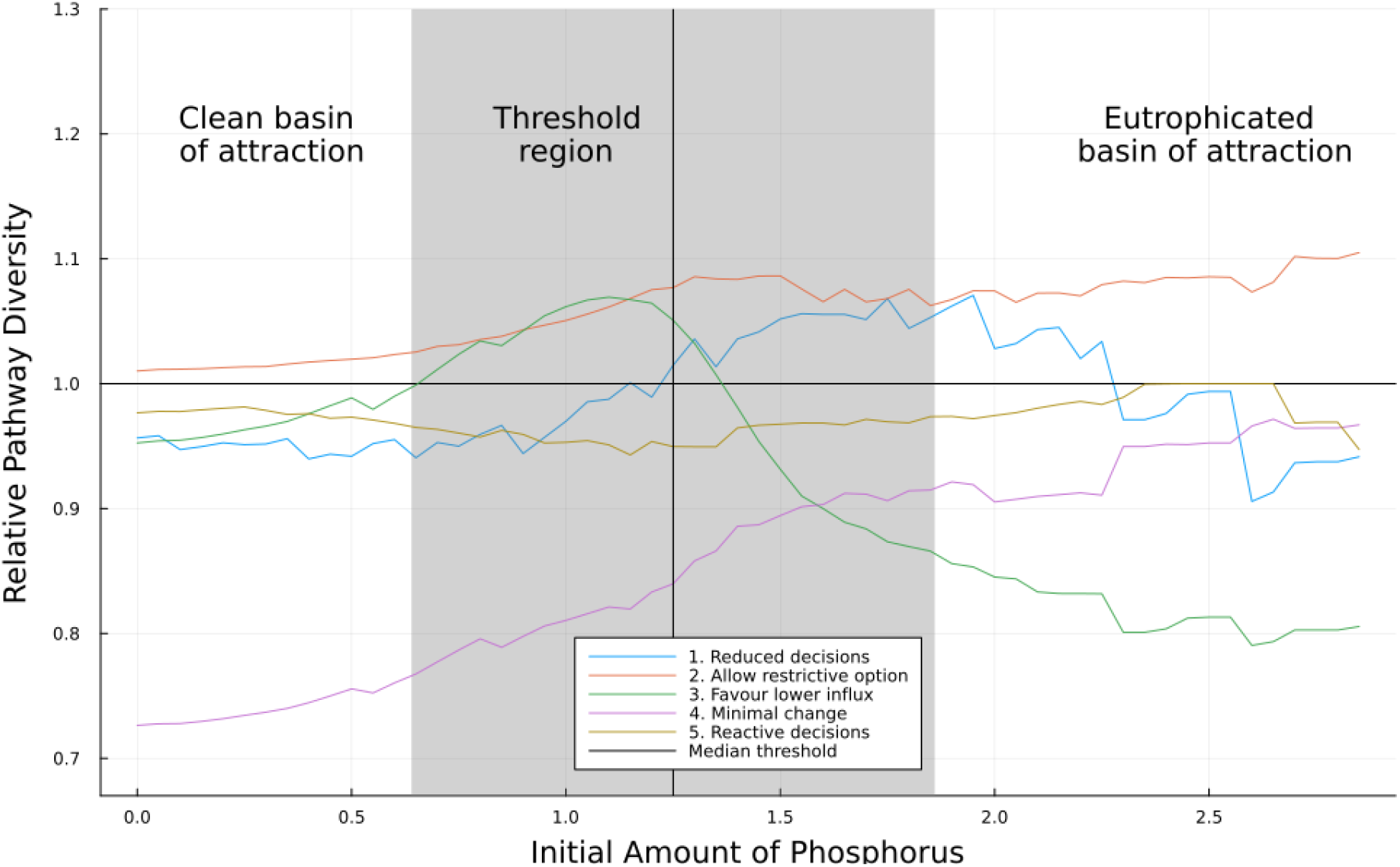
Relative pathway diversity over different initial conditions for five policy decision-making scenarios with different constraints on agency. All results are compared with the default scenario (*y=1*). For more details on the definitions of the scenarios, see Figure 3 and Table 1. The original data, without normalisation by the default scenario, is presented in Appendix S1: Figure S3.

Three scenarios have a decision-maker with lower capability and generally lower pathway diversity: scenarios 1, 4 and 5. In the *reduced decisions* scenario, the decision-maker has lower capability to adjust decisions and implement them within a given time horizon due to a high institutional rigidity. Reducing decision frequency lowers pathway diversity by limiting the number of pathways in most of the state space. Since decisions persist longer, a lower influx option just beyond the threshold can reverse the regime shift, leading to a higher pathway diversity in this region (Figure 7). In the *minimal change* scenario, the reduction in capability is caused by the decision-maker having the probability of choosing different options constrained by choosing options closer to the previously adopted option, introducing a strong path dependence that limits change. The scenario shows the lowest pathway diversity across most of the state space (Figure 7). The reduction in the number of options is higher in the clean basin of attraction, as is the decrease in pathway diversity. In the *reactive decisions* scenario, a crisis constrains the decision-maker’s capability and drives them into choosing short-term, reactive options based on past states. Pathway diversity decreases due to restrictions in the decision space, as the decision-maker consistently favours either high or low influx (Figure 7). This effect is slightly weaker past the basin threshold, where the state tends to rise, leading to lower influx choices and a higher chance of reversibility.

Two scenarios have a decision-maker with higher capability and generally higher pathway diversity: scenarios 2 and 3. In the *allow restrictive option* scenario, the increased capability is caused by a higher government legitimacy, which allow the enforcement of a considerably lower influx option. When following this new option, the system can enter a configuration where only the clean state is stable (Figure 1). Therefore, the system can revert the regime shift and return to the clean basin of attraction. This scenario produces a higher pathway diversity, especially in the eutrophic region (Figure 7). In the *favour lower influx* scenario, the decision-maker has greater capability to make and implement environmentally focused decisions due to an alignment between the decision-maker and community values. Favouring part of the decision space generally reduces pathway diversity. However, in this scenario, it increases pathway diversity around the threshold, as prioritising lower influx helps keep the system in the clean basin of attraction, where more options are available (Figure 7).

## 4. Discussion

Ecological resilience metrics have traditionally emphasised systemic elements and often miss the role of agency (Cote and Nightingale, 2012; Olsson et al., 2015; Sanches et al., 2025). In contrast, social-ecological resilience assessments often allow more nuance by incorporating both system and agent dimensions, though these assessments are typically qualitative (González-Quintero and Avila-Foucat, 2019; Polain de Waroux et al., 2024). This study highlights that pathway diversity provides a quantitative approach capable of capturing both dimensions while remaining compatible with previous approaches. Using a lake eutrophication model, we showed that the most significant decline in pathway diversity occurs within the threshold region (Figure 4) and that a pathway diversity time series can serve as a reliable early-warning signal (Figure 6). Furthermore, by using policy decision-making scenarios that constrain or enable the decision-maker’s capability to choose different options, we show that higher pathway diversity is associated with greater agency and decision-maker’ capability, though this effect can vary across the state space (Figure 7). This result illustrates how agency can shape system dynamics, influence the likelihood of regime shifts, and affect resilience.

We expect the results showing pathway diversity’s compatibility with ecological resilience metrics to be generalisable to a wide range of problems that have a clear link between decision and state space. In the lake eutrophication model, we assume that as the lake becomes eutrophic — change in state space — the decision-maker faces fewer influx options and must adopt stricter policies — change in decision space. While not essential for applying pathway diversity, this link is crucial for our observed compatibility result. It allows pathway diversity, which operates within the decision space, to capture the system’s dynamic and trajectory in the state space (see Figures 4–5 for further details). Drawing on normal form theory, which simplifies complex system behaviour near critical points by eliminating higher-order terms (Kuznetsov et al., 1998), we expect our results in the vicinity of the threshold to extend to more complex models if they exhibit bi-stability and each basin of attraction offers a different set of options. Testing pathway diversity in case studies with varying system complexity could validate this expectation. The link between state and decision space is common in social-ecological systems but can be more complex than in this study. For example, in the Gordon-Schaefer fisheries model (Gordon, 1954), the availability of catch limit options can depend on both ecological factors (e.g., fish population) and socio-economic factors (e.g., market demand). Building on the connection between decision and state space, integrating pathway diversity with approaches that explore state trajectories to measure resilience, such as ecological dynamic regimes (Sánchez-Pinillos et al., 2024) and viability theory (Béné and Doyen, 2018; Rougé et al., 2013), could enhance its applicability across different contexts.

Analyses of pathway diversity have the potential to identify implications for resilience-based policy (Garmestani and Benson, 2013), as it can capture how social-ecological resilience is shaped by interactions between biophysical dynamics and constraints on agency, as demonstrated in the scenarios here. Our model incorporates how internal and external factors affect the decision-maker’s agency in terms of their capability to choose different options. Two central concepts from the resilience literature can help understand the factors impacting agency and resilience: rigidity traps and biophysical lock-ins. Rigid policies and institutions that impose fixed rules or restrict choices to optimal options can lead to rigidity traps that undermine resilience by reducing pathway diversity, reinforcing the importance of considering multiple plausible options (Carpenter et al., 2015; Garmestani and Benson, 2013; Haasnoot et al., 2013; Moore and Milkoreit, 2020). Similarly, external pressures that reduce flexibility often increase path dependence and push systems into rigidity traps⍰where transformation becomes increasingly unlikely(Carpenter and Brock, 2008; Enqvist et al., 2016; Méndez et al., 2022). Meanwhile, decisions that increase the probability of entering a biophysical lock-in strongly reduce adaptive capacity and increase the probability of becoming trapped in undesirable conditions (Carpenter and Brock, 2008; Lade et al., 2017), where pathway diversity is low. These factors can interact, either supporting or limiting resilience. For example, the *favour lower influx* scenario increases decision rigidity but reduces biophysical lock-ins, resulting in a higher resilience in parts of the state space (Figure 7). This result illustrates that the effect on resilience can be state-dependent, such that the impact of a constraint on agency may be more complex than initially expected. Overall, our findings highlight the need to strengthen actors’ capability and provide a method to assess when these complex factors can improve resilience.

We suggest three future research directions to build on our results and advance the application of pathway diversity in decision-making. First, pathway diversity should be applied to more realistic decision-making models. While our model shows that pathway diversity can capture constraints on agency, it was not designed to reflect the full complexity of real-world decision processes. Implementing pathway diversity in more realistic models is important to strengthen tests of pathway diversity’s policy relevance. Achieving this goal will require advances in understanding and modelling individual human decision-making (Schlüter et al., 2012) so that we can identify how individual and system-level factors constrain agents’ options and conduct a meaningful pathway diversity analysis. Second, future work should assess whether there are limits to how much pathway diversity enhances resilience in terms of the capacity to deal with disturbances. In this study, higher pathway diversity is always associated with higher resilience, but in more complex models, as in reality, excessive diversity can increase system complexity and undermine resilience (Biggs et al., 2015). Ensuring options are defined at an appropriate level of granularity may help align pathway diversity with resilience. Third, further research is needed to clarify the link between pathway diversity and performance. Decision-making approaches typically select policies by optimising performance indicators based on economic welfare (Barfuss et al., 2018; Perman et al., 2003), while approaches like adaptive pathways emphasise flexibility while still considering performance (Haasnoot et al., 2013; Haasnoot et al., 2024). Pathway diversity can capture performance through feedback, and in our lake system, they are already closely linked — states with high pathway diversity have high performance (e.g., high provision of ecosystem services). Future research should analyse whether pathway diversity alone can produce meaningful policy recommendations or whether alternatives, such as multi-objective optimisation balancing resilience and performance, are necessary.

## 5. Conclusions

The recently proposed concept of pathway diversity offers a novel approach to analysing resilience, providing insights into how change in agency affects resilience. Using a lake eutrophication model and five policy decision-making scenarios, we demonstrate that pathway diversity captures changes in agency and is generally associated with bigger decision-makers’ capability of making decisions. At the same time, we show that pathway diversity is compatible with established ecological resilience metrics, capturing the risk of crossing a threshold, and serving as an early-warning signal. This study highlights the potential of pathway diversity to identify resilience-based policy implications, emphasising its value in integrating system and agency elements of resilience.

## 6. Acknowledgements

The research leading to these results has received funding from the Australian Government (Australian Research Council Future Fellowship FT200100381 to S.J.L.).

## 7. Author Contributions

VHS and SJL designed the study. VHS developed the software and wrote the first draft of the manuscript. JG, TI, SC, SJL contributed with supervision, discussion of the methodology and interpretation of the results. SJL was responsible for funding acquisition. All authors contributed with revision and editing of the manuscript.

## Appendix S1

**Figure S1:**
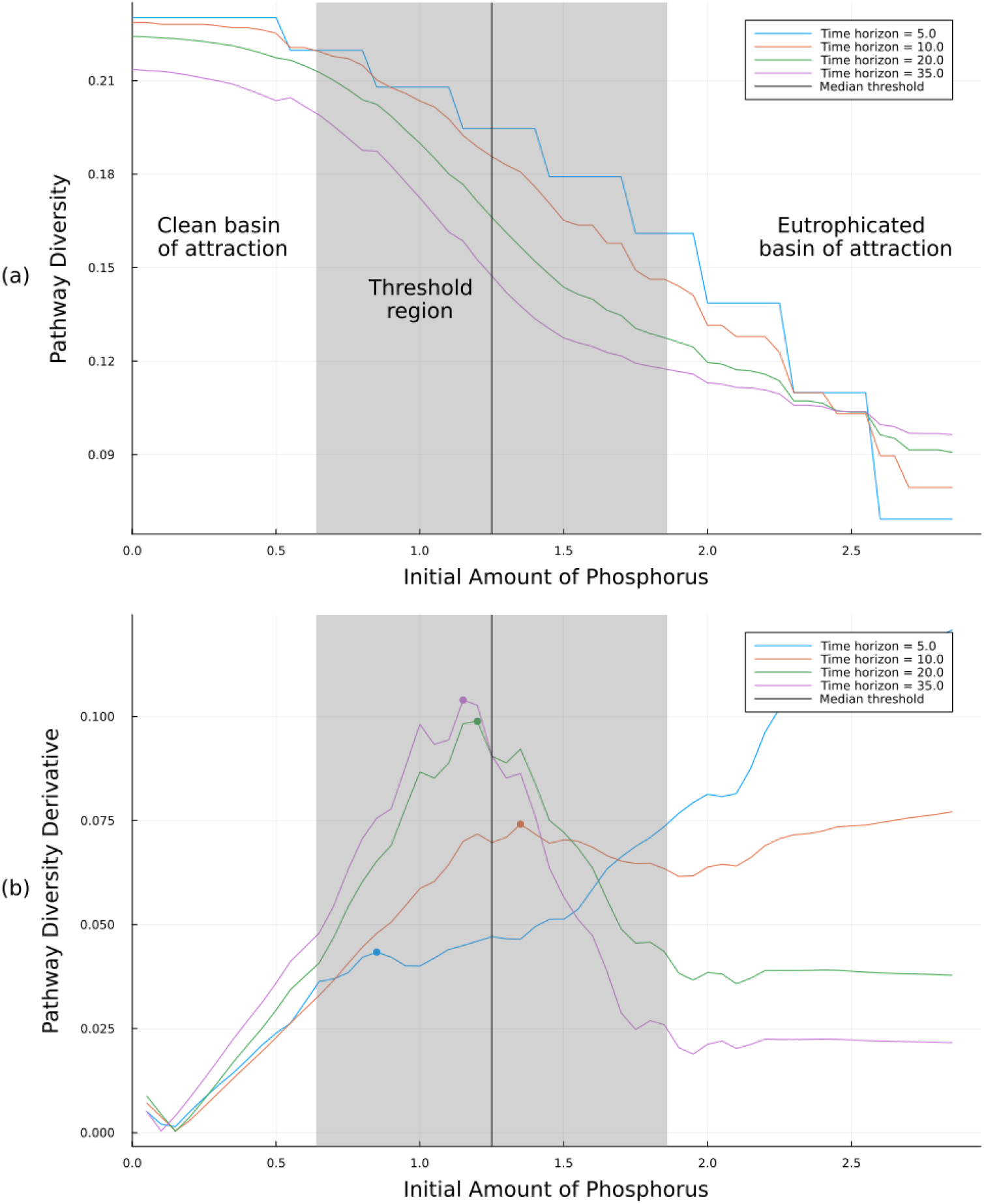
Pathway diversity for different time horizons and initial amounts of phosphorus (a) and the derivative of that curves (b). The threshold position varies - depending on the influx - within the range shown in grey and has a median indicated by the black vertical line. In (b), we show the derivative of a smoothed version of the pathway diversity. The points in (b) indicates the peak on the derivative, which happens in the threshold region. For the time-horizon of 5 years there is an increase on the pathway diversity derivative for higher initial amount of phosphorus due to discretisation problems.

**Figure S2:**
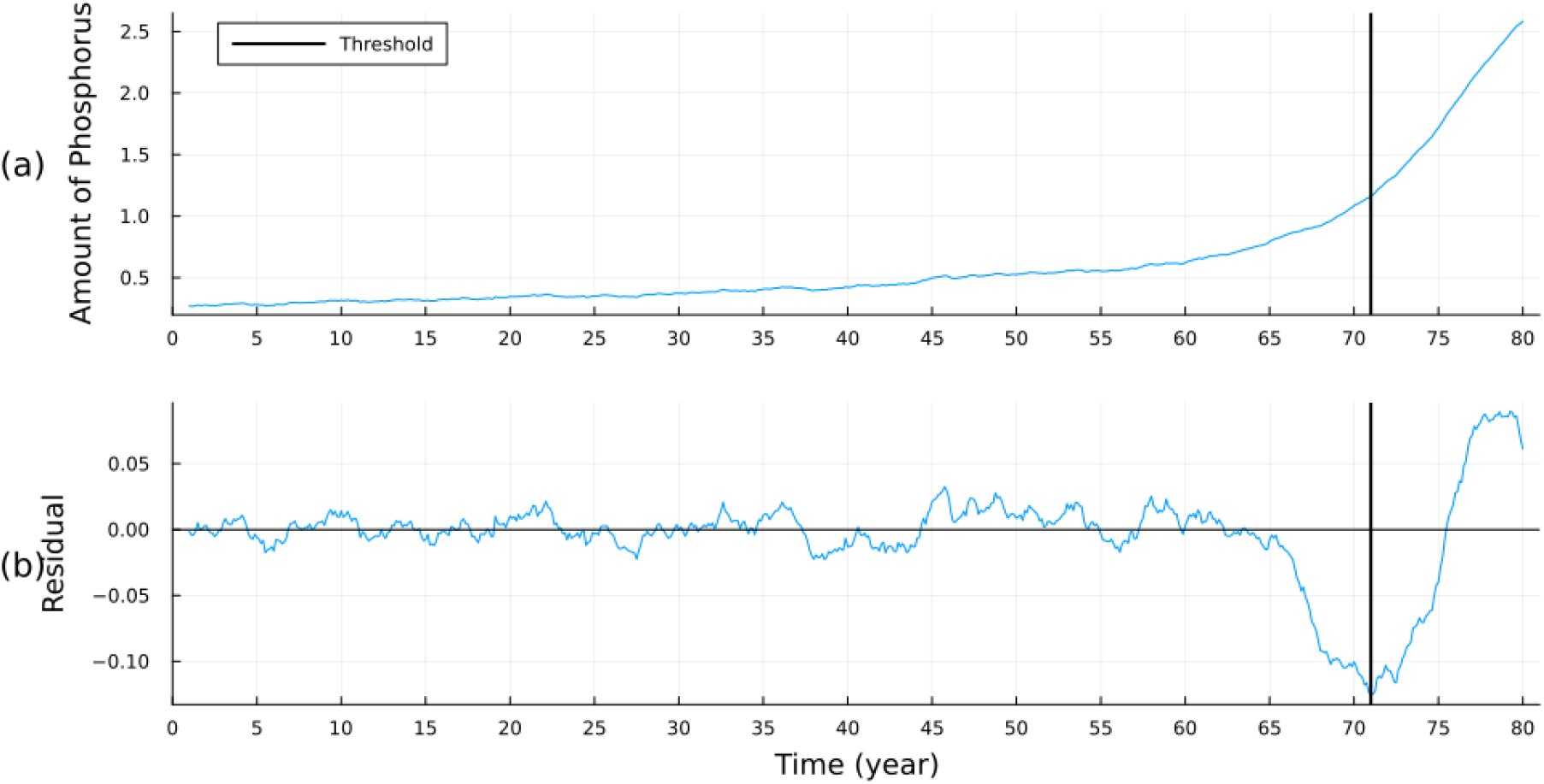
Time series of amount of phosphorus and its residual for the early-warning signal simulation. (a) Simulated amount of phosphorous (x), with a linear increase in the influx of phosphorus per time and white noise. (b) Residuals of the application of the LOESS detrending method. Note that around the year 65, the residuals start to deviate from zero.

**Figure S3:**
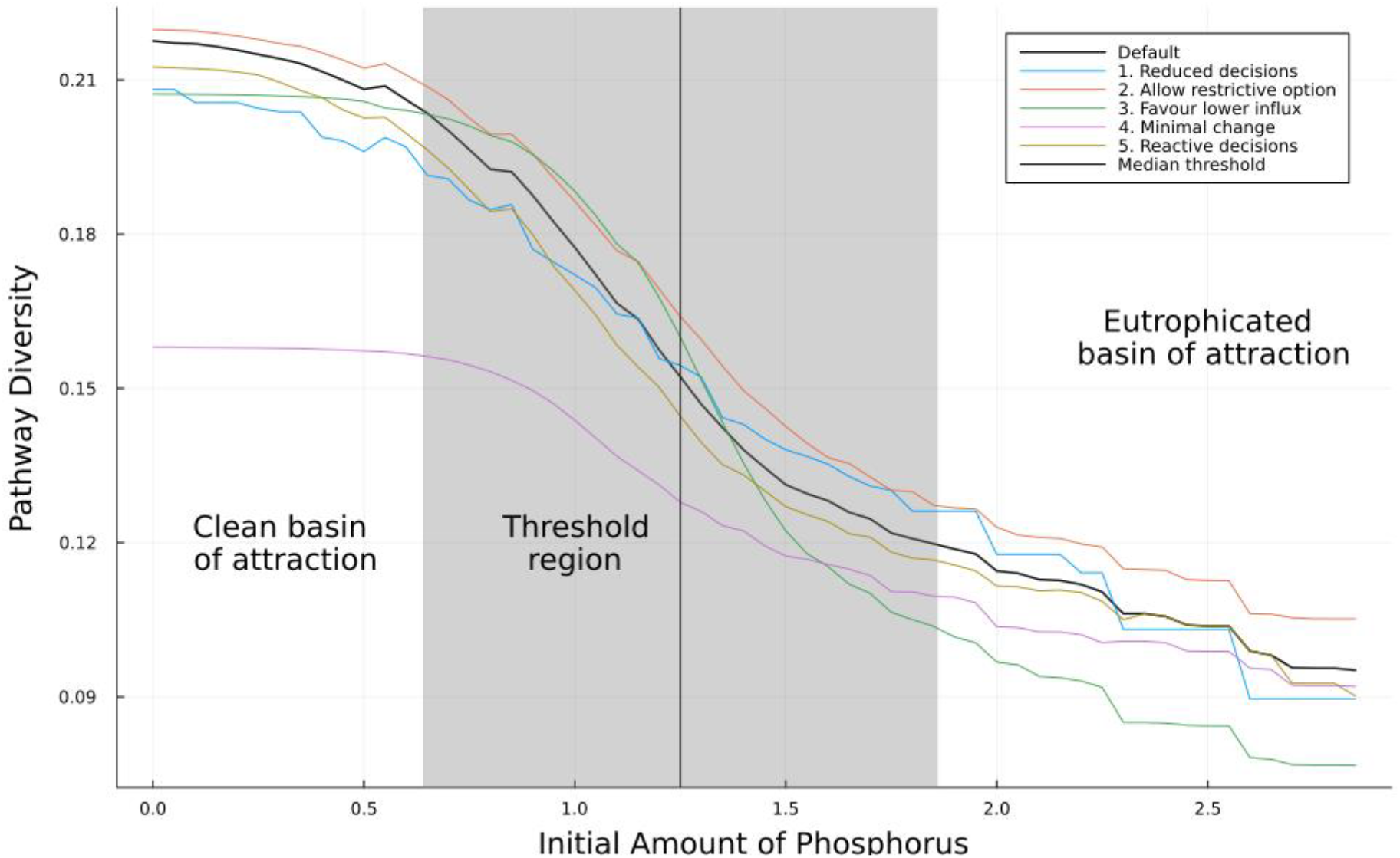
Pathway diversity over different initial conditions for five scenarios with different constraints on agency and a default scenario. For more details on the definitions of the scenarios, see Figure 3 and Table 1.

## References

Allen, C.R., Angeler, D.G., Chaffin, B.C., Twidwell, D., Garmestani, A., 2019. Resilience reconciled. Nature Sustainability 2, 898–900. 10.1038/s41893-019-0401-4

Baggio, J., Brown, K., Hellebrandt, D., 2015. Boundary object or bridging concept? A citation network analysis of resilience. Ecology and Society 20. 10.5751/ES-07484-200202

Barfuss, W., Donges, J.F., Lade, S.J., Kurths, J., 2018. When optimization for governing human-environment tipping elements is neither sustainable nor safe. Nature Communications 9, 2354. 10.1038/s41467-018-04738-z

Béné, C., Doyen, L., 2018. From Resistance to Transformation: A Generic Metric of Resilience Through Viability. Earth’s Future 6, 979–996. 10.1002/2017EF000660

Biggs, R., Peterson, G.D., Rocha, J.C., 2018. The Regime Shifts Database: a framework for analyzing regime shifts in social-ecological systems. Ecology and Society 23. 10.5751/ES-10264-230309

Biggs, R., Schlüter, M., Schoon, M.L., 2015. Principles for building resilience: sustaining ecosystem services in social-ecological systems. Cambridge University Press.

Bohle, H.-G., Etzold, B., Keck, M., 2009. Resilience As Agency. International Human Dimension Programme Update 2:8–13.

Brown, K., Westaway, E., 2011. Agency, Capacity, and Resilience to Environmental Change: Lessons from Human Development, Well-Being, and Disasters. Annual Review of Environment and Resources 36, 321–342. 10.1146/annurev-environ-052610-092905

Carpenter, S.R., Brock, W.A., 2008. Adaptive Capacity and Traps. Ecology and Society 13.

Carpenter, S.R., Brock, W.A., Folke, C., van Nes, E.H., Scheffer, M., 2015. Allowing variance may enlarge the safe operating space for exploited ecosystems. Proceedings of the National Academy of Sciences 112, 14384–14389. 10.1073/pnas.1511804112

Carpenter, S.R., Ludwig, D., Brock, W.A., 1999. Management of Eutrophication for Lakes Subject to Potentially Irreversible Change. Ecological Applications 9, 751–771. 10.1890/1051-0761(1999)009[0751:MOEFLS]2.0.CO;2

Cosens, B.A., 2013. Legitimacy, Adaptation, and Resilience in Ecosystem Management. Ecology and Society 18. 10.5751/ES-05093-180103

Cote, M., Nightingale, A.J., 2012. Resilience thinking meets social theory: Situating social change in socio-ecological systems (SES) research. Progress in Human Geography 36, 475–489. 10.1177/0309132511425708

Dakos, V., Carpenter, S.R., Brock, W.A., Ellison, A.M., Guttal, V., Ives, A.R., Kéfi, S., Livina, V., Seekell, D.A., Nes, E.H. van, Scheffer, M., 2012. Methods for Detecting Early Warnings of Critical Transitions in Time Series Illustrated Using Simulated Ecological Data. PLOS ONE 7, e41010. 10.1371/journal.pone.0041010

Dakos, V., Kéfi, S., 2022. Ecological resilience: what to measure and how. Environmental Research Letters 17, 043003. 10.1088/1748-9326/ac5767

Duncan, S., 2019. Agency, in: Atkinson, P., Delamont, S., Cernat, A., Sakshaug, J.W., Williams, R.A. (Eds.), SAGE Research Methods Foundations. SAGE Publications, In press. 10.4135/9781526421036811849

Elmqvist, T., Folke, C., Nyström, M., Peterson, G., Bengtsson, J., Walker, B., Norberg, J., 2003. Response diversity, ecosystem change, and resilience. Frontiers in Ecology and the Environment 1, 488–494. 10.1890/1540-9295(2003)001[0488:RDECAR]2.0.CO;2

Enqvist, J., Tengö, M., Boonstra, W.J., 2016. Against the current: rewiring rigidity trap dynamics in urban water governance through civic engagement. Sustain Sci 11, 919–933. 10.1007/s11625-016-0377-1

Folke, C., 2006. Resilience: The emergence of a perspective for social–ecological systems analyses. Global Environmental Change 16, 253–267. 10.1016/j.gloenvcha.2006.04.002

Garmestani, A.S., Benson, M.H., 2013. A Framework for Resilience-based Governance of Social-Ecological Systems. E&S 18, art9. 10.5751/ES-05180-180109

González-Quintero, C., Avila-Foucat, V.S., 2019. Operationalization and Measurement of Social-Ecological Resilience: A Systematic Review. Sustainability 11, 6073. 10.3390/su11216073

Gordon, H.S., 1954. The Economic Theory of a Common-Property Resource: The Fishery. Journal of Political Economy 62, 124–142.

Haasnoot, M., Di Fant, V., Kwakkel, J., Lawrence, J., 2024. Lessons from a decade of adaptive pathways studies for climate adaptation. Global Environmental Change 88, 102907. 10.1016/j.gloenvcha.2024.102907

Haasnoot, M., Kwakkel, J.H., Walker, W.E., ter Maat, J., 2013. Dynamic adaptive policy pathways: A method for crafting robust decisions for a deeply uncertain world. Global Environmental Change 23, 485–498. 10.1016/j.gloenvcha.2012.12.006

Hahn, T., Nykvist, B., 2017. Are adaptations self-organized, autonomous, and harmonious? Assessing the social-ecological resilience literature. Ecology and Society 22, art12. 10.5751/ES-09026-220112

Haider, L.J., Cleaver, F., 2023. Capacities for resilience: persisting, adapting and transforming through bricolage. Ecosystems and People 19, 2240434. 10.1080/26395916.2023.2240434

Holling, C.S., 1973. Resilience and Stability of Ecological Systems. Annual Review of Ecology and Systematics 4, 1–23. 10.1146/annurev.es.04.110173.000245

Jones, N.A., Shaw, S., Ross, H., Witt, K., Pinner, B., 2016. The study of human values in understanding and managing social-ecological systems. Ecology and Society 21. 10.5751/ES-07977-210115

Jozaei, J., Chuang, W.-C., Allen, C.R., Garmestani, A., 2022. Social vulnerability, social-ecological resilience and coastal governance. Global Sustainability 5, e12. 10.1017/sus.2022.10

Krakovská, H., Kuehn, C., Longo, I.P., 2023. Resilience of dynamical systems. European Journal of Applied Mathematics 1–46. 10.1017/S0956792523000141

Kuznetsov, Y.A., 1998. Elements of Applied Bifurcation Theory, Applied Mathematical Sciences. Springer, New York. 10.1007/978-3-031-22007-4

Lade, S.J., Haider, L.J., Engström, G., Schlüter, M., 2017. Resilience offers escape from trapped thinking on poverty alleviation. Science Advances 3, e1603043. 10.1126/sciadv.1603043

Lade, S.J., Walker, B.H., Haider, L.J., 2020. Resilience as pathway diversity: linking systems, individual, and temporal perspectives on resilience. Ecology and Society 25, art19. 10.5751/ES-11760-250319

Laurien, F., Martin, J.G.C., Mehryar, S., 2022. Climate and disaster resilience measurement: Persistent gaps in multiple hazards, methods, and practicability. Climate Risk Management 37, 100443. 10.1016/j.crm.2022.100443

Méndez, P.F., Fajardo-Ortiz, D., Holzer, J.M., 2022. Chapter Nine - Disrupting the governance of social-ecological rigidity traps: Can pluralism foster change towards sustainability?, in: Holzer, J.M., Baird, J., Hickey, G.M. (Eds.), Advances in Ecological Research, Pluralism in Ecosystem Governance. Academic Press, pp. 243–291. 10.1016/bs.aecr.2022.04.011

Moore, M.-L., Milkoreit, M., 2020. Imagination and transformations to sustainable and just futures. Elementa: Science of the Anthropocene 8, 081. 10.1525/elementa.2020.081

Morgan, M., Webster, A.J., Padowski, J.C., Morrison, R.R., Flint, C.G., Simmons-Potter, K., Chief, K., Litson, B., Neztsosie, B., Karanikola, V., Kacira, M., Rushforth, R.R., Boll, J., Stone, M.B., 2024. Guided transformations for communities facing social and ecological change. Ecology and Society 29. 10.5751/ES-15448-290420

Olsson, L., Jerneck, A., Thoren, H., Persson, J., O’Byrne, D., 2015. Why resilience is unappealing to social science: Theoretical and empirical investigations of the scientific use of resilience. Science Advances 1, e1400217. 10.1126/sciadv.1400217

Otto, I.M., Wiedermann, M., Cremades, R., Donges, J.F., Auer, C., Lucht, W., 2020. Human agency in the Anthropocene. Ecological Economics 167, 106463. 10.1016/j.ecolecon.2019.106463

Perman, R.J., Ma, Y., Common, M., Maddison, D., McGilvray, J.W., 2003. Natural resource and environmental economics. Pearson Education, Essex.

Peterson, G.D., Carpenter, S.R., Brock, W.A., 2003. Uncertainty and the Management of Multistate Ecosystems: An Apparently Rational Route to Collapse. Ecology 84, 1403–1411. 10.1890/0012-9658(2003)084[1403:UATMOM]2.0.CO;2

Polain de Waroux, Y., Carignan, M.-C., del Giorgio, O., Díaz, L., Enrico, L., Jaureguiberry, P., Lipoma, M.L., Mazzini, F., Díaz, S., 2024. How do we study resilience? A systematic review. People and Nature 6, 474–489. 10.1002/pan3.10603

Quinlan, A.E., Berbés-Blázquez, M., Haider, L.J., Peterson, G.D., 2016. Measuring and assessing resilience: broadening understanding through multiple disciplinary perspectives. Journal of Applied Ecology 53, 677–687. 10.1111/1365-2664.12550

Quinn, J.D., Reed, P.M., Keller, K., 2017. Direct policy search for robust multi-objective management of deeply uncertain socio-ecological tipping points. Environmental Modelling & Software 92, 125– 141. 10.1016/j.envsoft.2017.02.017

Reyers, B., Moore, M.-L., Haider, L.J., Schlüter, M., 2022. The contributions of resilience to reshaping sustainable development. Nat Sustain 5, 657–664. 10.1038/s41893-022-00889-6

Rougé, C., Mathias, J.-D., Deffuant, G., 2013. Extending the viability theory framework of resilience to uncertain dynamics, and application to lake eutrophication. Ecological Indicators 29, 420–433. 10.1016/j.ecolind.2012.12.032

Sanches, V., Quiñones, R., Vivas, J., Guillaume, J., Iwanaga, T., Kwakkel, J., Quinlan, A., Rocha, J., Crépin, A.-S., Dakos, V., Donges, J., Lade, S., 2025. Integrating Diversity and Agency into Social-Ecological Resilience Metrics. 10.31219/osf.io/2g3jp_v1

Sánchez-Pinillos, M., Dakos, V., Kéfi, S., 2024. Ecological dynamic regimes: A key concept for assessing ecological resilience. Biological Conservation 289, 110409. 10.1016/j.biocon.2023.110409

Scheffer, M., Bascompte, J., Brock, W.A., Brovkin, V., Carpenter, S.R., Dakos, V., Held, H., van Nes, E.H., Rietkerk, M., Sugihara, G., 2009. Early-warning signals for critical transitions. Nature 461, 53–59. 10.1038/nature08227

Scheffer, M., Carpenter, S., Foley, J.A., Folke, C., Walker, B., 2001. Catastrophic shifts in ecosystems. Nature 413, 591–596. 10.1038/35098000

Schlüter, M., Mcallister, R.R.J., Arlinghaus, R., Bunnefeld, N., Eisenack, K., Hölker, F., Milner-Gulland, .j., Müller, B., Nicholson, E., Quaas, M., Stöven, M., 2012. New Horizons for Managing the Environment: A Review of Coupled Social-Ecological Systems Modeling. Natural Resource Modeling 25, 219–272. 10.1111/j.1939-7445.2011.00108.x

Sharifi, A., 2016. A critical review of selected tools for assessing community resilience. Ecological Indicators 69, 629–647. 10.1016/j.ecolind.2016.05.023

Singh, R., Reed, P., Keller, K., 2015. Many-objective robust decision making for managing an ecosystem with a deeply uncertain threshold response. Ecology and Society 20. 10.5751/ES-07687-200312

Steinmann, P., Tobi, H., van Voorn, G.A.K., 2024. Resilience Metrics for Socio-Ecological and Socio-Technical Systems: A Scoping Review. Systems 12, 357. 10.3390/systems12090357

Walker, B., Crépin, A.-S., Nyström, M., Anderies, J.M., Andersson, E., Elmqvist, T., Queiroz, C., Barrett, S., Bennett, E., Cardenas, J.C., Carpenter, S.R., Chapin, F.S., de Zeeuw, A., Fischer, J., Folke, C., Levin, S., Nyborg, K., Polasky, S., Segerson, K., Seto, K.C., Scheffer, M., Shogren, J.F., Tavoni, A., van den Bergh, J., Weber, E.U., Vincent, J.R., 2023. Response diversity as a sustainability strategy. Nature Sustainability 1–9. 10.1038/s41893-022-01048-7

Walker, B., Holling, C.S., Carpenter, S., Kinzig, A., 2004. Resilience, adaptability and transformability in social–ecological systems. Ecology and society 9.

Westley, F.R., Tjornbo, O., Schultz, L., Olsson, P., Folke, C., Crona, B., Bodin, Ö., 2013. A Theory of Transformative Agency in Linked Social-Ecological Systems. Ecology and Society 18. 10.5751/ES-05072-180327

Wissner-Gross, A.D., Freer, C.E., 2013. Causal Entropic Forces. Physical Review Letters 110, 168702. 10.1103/PhysRevLett.110.168702

